## Supplemental Figure for "A Host Enzyme Reduces Metabolic Dysfunction-Associated Steatotic Liver Disease (MASLD) by Inactivating Intestinal Lipopolysaccharide"

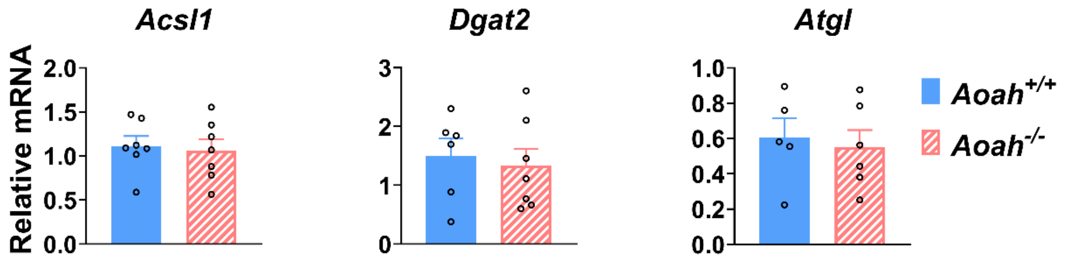


**Fig S1.** ***Aoah*$^{+/+}$ and *Aoah*$^{-/-}$ mouse livers express similar levels of triacylglycerol metabolism mRNA**

ACSL1, DGAT2 and ATGL mRNA was measured in co-housed 6 – 8 weeks old *Aoah*$^{+/+}$ and *Aoah*$^{-/-}$ mouse livers， n = 5 – 7.

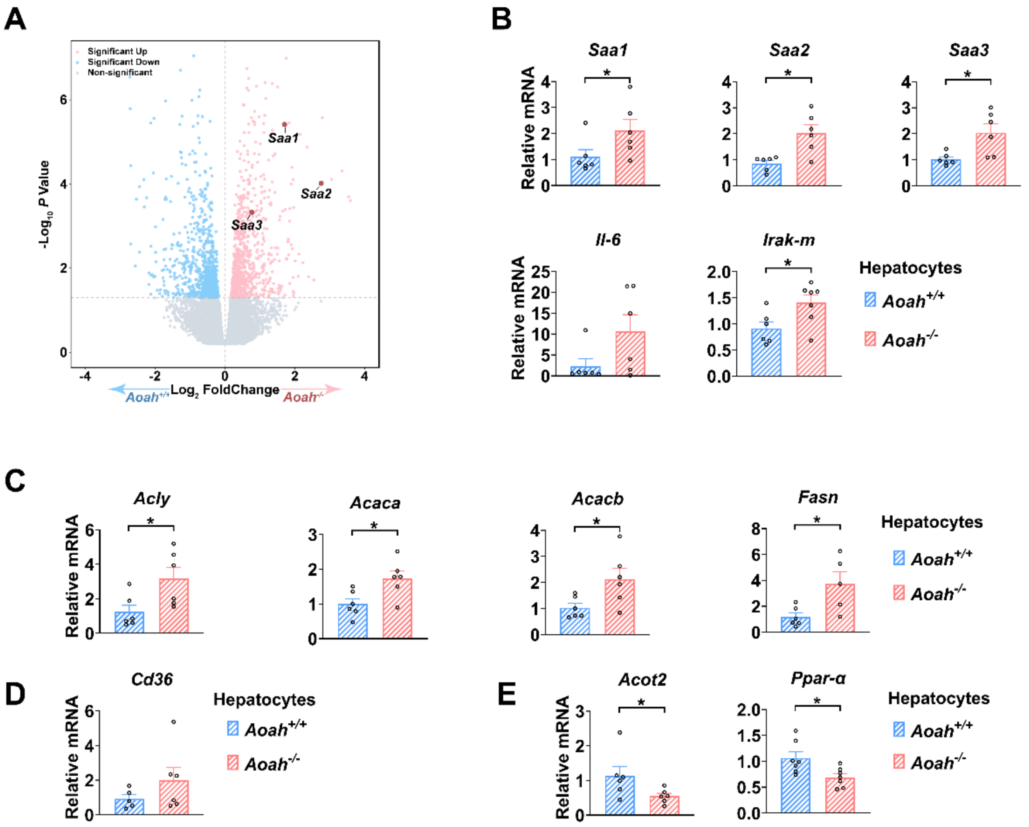


**Fig S2.** ***Aoah$^{-/-}$*** **mouse hepatocytes have altered expression of genes that may promote lipid storage.**

(A) The differentially expressed genes between livers from co-housed 6 – 8 weeks old *Aoah$^{+/+}$* and *Aoah$^{-/-}$* mouse are shown.

(B-E) Hepatocytes were isolated from co-housed 6 – 8 weeks old *Aoah$^{+/+}$* and *Aoah$^{-/-}$* mice and cell lysate was used for qPCR analysis for inflammatory (B) and fatty acid synthesis (C), uptake (D) and oxidation (E) gene expression. Data were combined from at least 2 experiments, n = 5 – 7.

Mann-Whitney test was used. *, $P < 0.05$.

### Table S1. Primers used for qPCR

| Mouse gene symbols | Forward primer sequence | Reverse primer sequence |
|---|---|---|
| *Actin* | 5'-GGCTGTATTCCCCTCCATCG-3' | 5'-CCAGTTGGTAACAATGCCATGT-3' |
| *Il-6* | 5'-ATCGTGGAAATGAGAAAAGAGTTGT-3' | 5'-AAGTGCATCATCGTTGTTCATACA-3' |
| *Tnf-α* | 5'-CATCTTCTCAAAATTCGAGTGACAA-3' | 5'- TCAGCCACTCCAGCTGCTC-3' |
| *Ifn-γ* | 5'-ATGAACGCTACACACTGCATC-3' | 5'-CCATCCTTTTGCCAGTTCCTC-3' |
| *Il-10* | 5'-GCTGGACAACATACTGCTAACC-3' | 5'-ATTTCCGATAAGGCTTGGCAA-3' |
| *Timp1* | 5'-GCAACTCGGACCTGGTCATAA-3' | 5'-CGGCCCGTGATGAGAAACT-3' |
| *Mmp2* | 5'-CAAGTTCCCCGGCGATGTC-3' | 5'-TTCTGGTCAAGGTCACCTGTC-3' |
| *α-SMA* | 5'-TCCACCGCAAATGCTTCTAAG-3' | 5'-TGTTGCTAGGCCAGGGCTAC-3' |
| *Collagen 4α1* | 5'-CTGGCACAAAAGGGACGAG-3' | 5'-ACGTGGCCGAGAATTTCACC-3' |
| *Cd36* | 5'-ATGGGCTGTGATCGGAACTG-3' | 5'-GTCTTCCCAATAAGCATGTCTCC-3' |
| *Fabp3* | 5'-ACCTGGAAGCTAGTGGACAG-3' | 5'-TGATGGTAGTAGGCTTGGTCAT-3' |
| *Fasn* | 5'-GGAGGTGGTGATAGCCGGTAT-3' | 5'-TGGGTAATCCATAGAGCCCAG-3' |
| *Acsl1* | 5'-TGCCAGAGCTGATTGACATTC-3' | 5'-GGCATACCAGAAGGTGGTGAG-3' |
| *Dgat2* | 5'-GCGCTACTTCCGAGACTACTT-3' | 5'-GGGCCTTATGCCAGGAAACT-3' |
| *Atgl* | 5'-GGATGGCGGCATTTCAGACA-3' | 5'-CAAAGGGTTGGGTTGGTTCAG-3' |
| *Pcx* | 5'-CTGAAGTTCCAAACAGTTCGAGG-3' | 5'-CGCACGAAACACTCGGATG-3' |
| *Acly* | 5'-AATCCTGGCTAAAACCTCGCC-3' | 5'-GCATAGATGCACACGTAGAACT-3' |
| *Acaca* | 5'-ATGGGCGGAATGG TCTCTTTC-3' | 5'-TGGGGACCTTGTCTTCATCAT-3' |
| *Acacb* | 5'-CCTTTGGCAACAAGCAAGGTA-3' | 5'-AGTCGTACACATAGGTGGTCC-3' |

| | | |
|---|---|---|
| *Scd1* | 5’-GCCAGACCGGGCTGAACACC-3’ | 5’-GGCCTCCCAAGTGCAGCAGG-3’ |
| *Acss2* | 5’-AAACACGCTCAGGGAAAATCA-3’ | 5’-ACCGTAGATGTATCCCCCAGG-3’ |
| *Acot2* | 5’-CCCCAAGAGCATAGAAACCA-3’ | 5’-CCAATTCCAGGTCCTTTTACC-3’ |
| *Ppar-α* | 5’-AGAGCCCCATCTGTCCTCTC-3’ | 5’-ACTGGTAGTCTGCAAAACCAAA-3’ |
| *Aoah* | 5’-CAGCTACTCCCATGGCCAAA-3’ | 5’-GCCACCTGGACTGAAGAGTT-3’ |
| *Srebf1a* | 5’-GATGTGCGAACTGGACACAGC-3’ | 5’-GAGAAGCTCTCAGGAGAGTTGG-3’ |
| *Srebf1c* | 5’-CGCGGACCACGGAGCCATG-3’ | 5’-GAGAAGCTCTCAGGAGAGTTGG-3’ |
| *Saa1* | 5’-TTGTTCACGAGGCTTTCC-3’ | 5’-TGAGCAGCATCATAGTTCC-3’ |
| *Saa2* | 5’-TGGCTGGAAAGATGGAGACAA-3’ | 5’-AAAGCTCTCTCTTGCATCACTG-3’ |
| *Saa3* | 5’-TGCCATCATTCTTTGCATCTTGA-3’ | 5’-CCGTGAACTTCTGAACAGCCT-3’ |
| *Irak-m* | 5’-TCCCACCTGAGGTGAAGCAT-3’ | 5’-TGTGACATTGGCTGGTTCCA-3’ |
